# Building a virtual summer research experience in cancer for high school and early undergraduate students: lessons from the COVID-19 pandemic

**DOI:** 10.1101/2020.11.23.393967

**Authors:** Timothy W Corson, Shannon M Hawkins, Elmer Sanders, Jessica Byram, Leigh-Ann Cruz, Jacob Olson, Emily Speidell, Rose Schnabel, Adhitya Balaji, Osas Ogbeide, Julie Dinh, Amy Hinshaw, Laura Cummings, Vicki Bonds, Harikrishna Nakshatri

**Affiliations:** Indiana University Melvin and Bren Simon Comprehensive Cancer Center, Indiana University, Indianapolis, IN 46202, USA; Department of Ophthalmology, Indiana University School of Medicine, Indianapolis, IN 46202, USA; Department of Obstetrics and Gynecology, Indiana University School of Medicine, Indianapolis, IN 46202, USA; Indiana Clinical and Translational Sciences Institute, Indianapolis, IN 46202, USA; K-12 STEM Program, Indiana University School of Medicine, IN 46202, USA; Department of Anatomy, Cell Biology and Physiology, Indiana University School of Medicine, IN 46202, USA; Riverside High School, Indianapolis, IN 46208, USA; Decatur Central High School, Indianapolis, IN 46221, USA; Indiana University, Bloomington, IN 47405, USA; Lawrence Township Schools, Indianapolis, IN 46226, USA; Herron High School, Indianapolis, IN 46202, USA; Pipeline and Pre-doctoral program, Indiana University School of Medicine, IN 46202, USA; Department of Surgery, Indiana University School of Medicine, Indianapolis, IN 46202, USA; Department of Biochemistry and Molecular Biology, Indianapolis, IN 46202, USA; VA Roudebush Medical Center, Indianapolis, IN 46202, USA

**Keywords:** research education, mentoring, virtual education, pipeline program, curriculum development

## Abstract

**Background:** The COVID-19 pandemic posed a unique challenge for summer research programs in 2020, particularly for programs aimed at hands-on experience for younger trainees. The Indiana University Melvin and Bren Simon Comprehensive Cancer Center supports two pipeline programs, which traditionally immerse high school juniors, seniors, and early undergraduate students from underrepresented populations in science in hands-on projects in cancer biology labs. However, due to social distancing policies during the pandemic and reduction of research operations, these students were not physically allowed on campus. Thus, the authors set out to strategically pivot to a wholly virtual curriculum and evaluate the Virtual Summer Research Experience in Cancer outcomes.

**Methods:** The virtual program included four components: 1. a core science and professional development curriculum led by high school teachers and senior undergraduates; 2. faculty-delivered didactic sessions on cancer science; 3. mentored, virtual research projects with research faculty; and 4. online networking events to encourage vertical mentoring. Outcomes data were measured using an 11-item Research Preparation scale, daily electronic feedback, and structured evaluation and feedback via Zoom weekly.

**Results:** Outcome data suggested high self-reported satisfaction with the virtual program. Outcome data also revealed the importance of coordination between multiple entities for seamless program implementation. This includes the active recruitment and participation of high school teachers and further investment in information technology capabilities of institutions.

**Conclusions:** Findings reveal a path to educate and train high school and early undergraduate students in cancer research when hands-on, in-person training is not feasible. Virtual research experiences are not only useful to engage students during public health crises but can provide an avenue for cancer centers to expand their cancer education footprints to remotely located schools and universities with limited resources to provide such experiences to their students.

## Background

The COVID-19 pandemic resulted in significant challenges to the United States healthcare system [1, 2]. Many Academic Health Centers and Cancer Centers were charged with maintaining the traditional tripartite mission of clinical care, research, and education. Within medical education, virtual or distance learning combined with simulation became more common [1, 3]. Similarly, graduate medical and research education incorporated virtual didactic and telemedicine training [3–7]. Here, we describe the strategic pivot to the Virtual Summer Research Experience in Cancer (vSREC) from two traditional pipeline programs, aimed at immersing high school juniors, seniors, and early undergraduate students from underrepresented populations in biomedical science.

Providing early biomedical research opportunities has been shown to enhance future interest in biomedical careers [8]. Student-reported gains included disciplinary skills, research design, information or data analysis skills, information literacy, self-confidence, communication, and professional advancement [9–11]. Importantly, the impact of early biomedical research experiences is higher among students from underrepresented backgrounds [12, 13]. These research experiences and the resulting sense of responsibility positively impact academic and career success after accounting for parental income, IQ, and other factors that influence achievement [14]. In addition to focusing on diverse student trainees, teachers’ participation in research programs that include laboratory research and professional development can improve their students’ achievement in science [15].

Several National Cancer Institute (NCI)-designated cancer centers have instituted summer research programs (SRP) for high school and early undergraduate students underrepresented in biomedical research. Since 2003, the Indiana University Simon Comprehensive Cancer Center (IUSCCC) has provided summer research experiences to over 300 students, hereafter termed interns, from underrepresented populations, defined using the NIH definition of populations underrepresented in the extramural biomedical workforce (detailed in Additional file 1). In addition, in 2013, IUSCCC launched the Future Scientist Program (FSP), focusing on high school juniors in the Indianapolis Public School district, which contains a high percentage of disadvantaged students. The two-month-long programs not only provided first-hand research experience in cancer but also allowed students to develop long-term professional relationships with faculty mentors. Over 70% of interns have entered healthcare/science professions, and several have become physician-scientists, physicians, or biomedical scientists (unpublished data).

Previously, SRP and FSP had a similar structure: interns received a stipend to work on a research project in a faculty mentor’s laboratory (usually bench-based research) for 6–8 weeks, culminating in a poster and/or oral presentation. The laboratory experience was enriched by attendance at guest lectures on cancer biology and clinical cancer care, workshops on college/medical/graduate school applications and professional etiquette, and formal didactic training in research ethics, responsible conduct of research, and use of animals in research. Also, interns had social and celebratory events along with vertical mentoring opportunities with other trainees to teach how to network and navigate the university environment. IUSCCC also more recently initiated a 3–4-week high school teacher research program (TRP), placing teachers in research laboratories for hands-on experience.

Preparation to launch SRP, FSP, and TRP for the 2020 summer started in Fall 2019 (Figure 1A), and application review, interviews, and candidate selection were almost complete just before the COVID-19 pandemic caused by the SARS-CoV2 novel coronavirus [16] forced a “hibernation” of research on our campus, pausing all but essential in-person research, as Indiana and much of the United States were placed under stay-at-home orders [17]. Since an in-person program became impossible, we opted to retool the curriculum as a virtual experience because of the importance of the programs in the lives of young interns, not just as a career-enhancing experience, but also as a full-time, stipend-based activity in a summer with few other options.

**Fig. 1.**
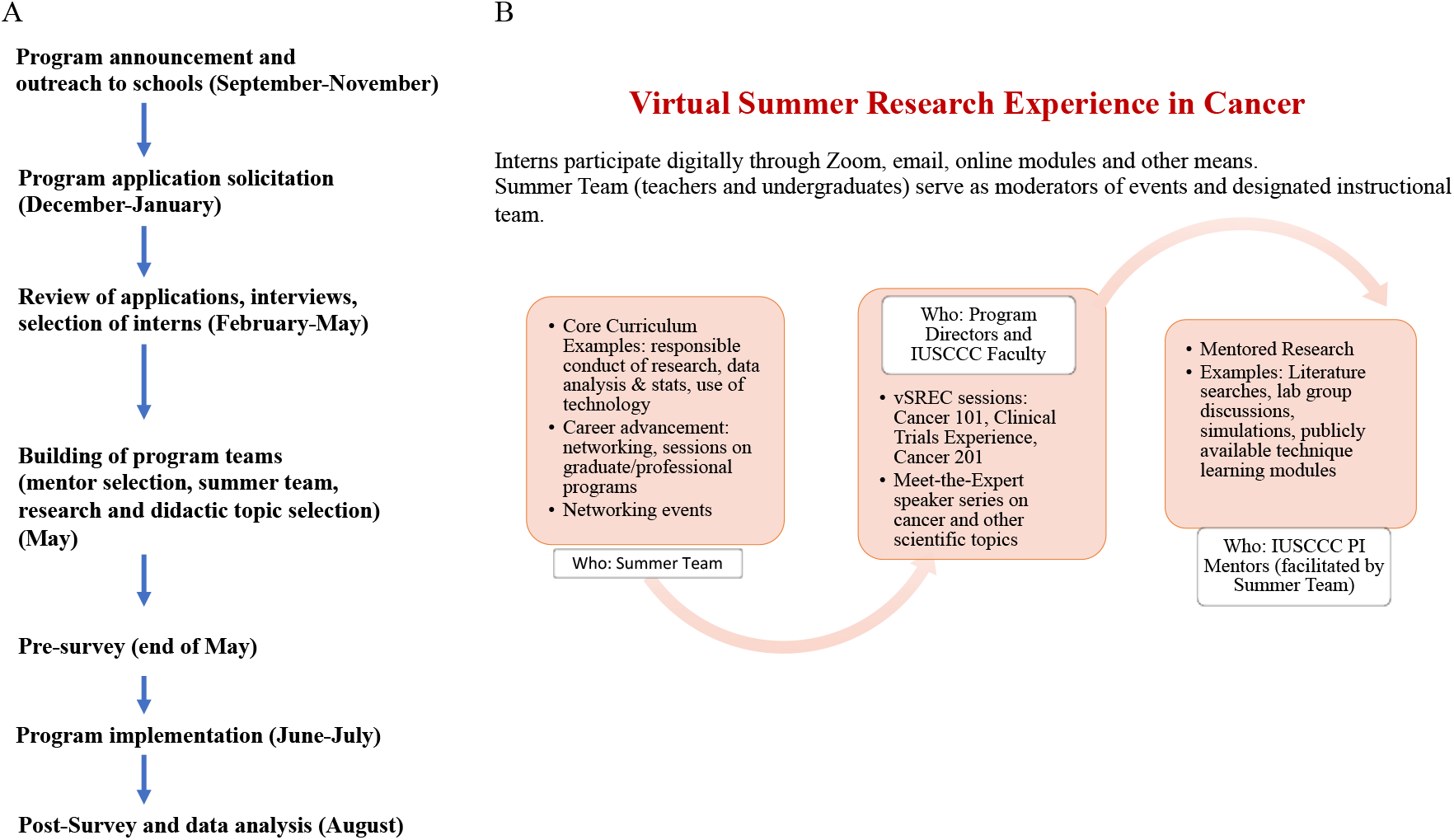
Schematic presentation of vSREC. A) Timeline and workflow of vSREC. B) vSREC core curriculum and participating teams and Faculty.

Here, we describe how the traditional SRP and FSP pipeline programs were modified into a virtual summer program, named Virtual Summer Research Experience in Cancer (vSREC), and its impact on participating interns. This first-of-its-kind virtual pipeline program is unique in that it brought together a diverse group of high school and undergraduate students, high school teachers, IUSCCC leadership, and faculty mentors with a shared goal to provide a positive experience in early biomedical research. Such a program is not only useful in future situations that require virtual learning, but also could be implemented on a routine basis to provide summer opportunities to students from rural school districts, non-research-intensive universities, or universities not affiliated with a cancer center or medical school.

## Methods

### Study Participants

All vSREC interns were invited to participate from May 2020 to July 2020. Intern demographics are detailed in Table 1 and included six Caucasian, 14 African American, one Asian, and one mixed-race interns. This study received approval from the Institutional Review Board at Indiana University (IRB protocol #1110007280). All procedures followed were in accordance with the ethical standards of the responsible committee on human experimentation and with the declaration of Helsinki.

**Table 1.** Intern demographics. n=22 interns total.

| Characteristic | n (%) |
| --- | --- |
| Gender |  |
| Male | 4 (18%) |
| Female | 18 (82%) |
| Educational level completed |  |
| High school junior | 8 (36%) |
| High school senior | 5 (23%) |
| College freshman | 9 (41%) |
| Race |  |
| Caucasian | 6 (27%) |
| African-American | 14 (64%) |
| Asian | 1 (5%) |
| Multi-Racial | 1 (5%) |
| Ethnicity |  |
| Hispanic/Latinx | 3 (14%) |
| Non-Hispanic/Latinx | 19 (86%) |

### Design of the vSREC

Figures 1A and B provide a schematic timeline and overview of the vSREC; additional details on program design are in Additional file 2. The educational objectives of vSREC mirrored the objectives of the in-person programs of previous years: to expose interns to university-level cancer research through a full-time, paid summer program; to introduce concepts in cancer biology and medicine; to inspire interns to pursue further studies in science and/or medicine; and to build long-term relationships between mentors and interns. For vSREC, we added the objective of enhancing contemporary scientific literacy through education on virology and SARS-CoV2, as new knowledge in this area was moving incredibly rapidly in the summer of 2020, along with significant dissemination of misinformation [18]. Additionally, we aimed to provide hands-on training for dealing with diversity and inclusion. This training was particularly timely during the Summer 2020 period of Black Lives Matter protests across the United States and recognition of racism, discrimination, and microaggressions in health care settings [19–22].

To meet all these objectives, vSREC utilized input and expertise in technology, teaching and evaluation, and cancer biology from a diverse group of individuals. Technology-adept local undergraduate students, who had completed rigorous science coursework, had a desire to pursue health- or science-related fields, had laboratory research experience, and/or had completed previous summer research programs on campus, provided hands-on and competent technology support, vertical mentoring, and campus navigation and networking advice. Local science teachers designed and delivered a 6-week core curriculum covering topics related to the research processes, scientific literacy, ethics, and grade-level resources for academic and career advancement. IUSCCC faculty delivered engaging lectures in cancer biology, starting with fundamental cancer topics and moving through areas of specialty, while also modeling various career paths. Faculty also served as research project mentors along with their laboratory groups, providing virtual projects that could be done remotely. These included in silico analyses, literature reviews, and analyses of existing imaging or other datasets, plus virtual training in laboratory techniques. Finally, engaging networking events gave students the chance to interact with peers and others. Further details of these components are provided in Additional file 2. An example intern’s weekly program schedule and activities are depicted in Figure 2, and Additional file 3 details all curriculum events, the daily checkout questions, and the extensive list of questions interns posed during the closing Cancer 201 lecture.

**Fig. 2.**
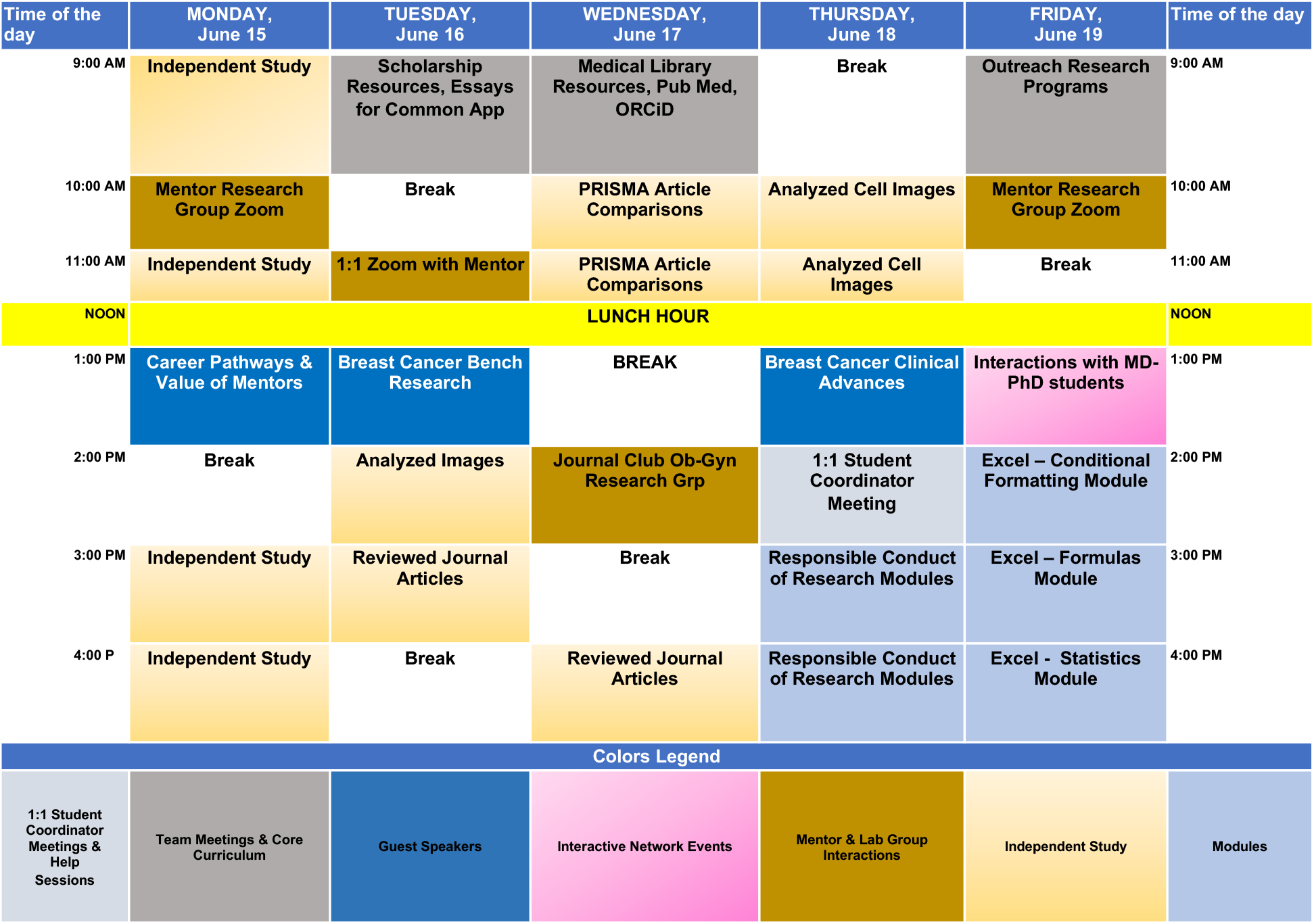
Representative vSREC weekly activity schedule. Activity of week 2 of the program for a specific intern is shown.

### Evaluation Methods

Survey instruments were created locally to evaluate the interns’ perceptions of the vSREC educational and research experience, their mentors, and the skills learned from the virtual experience. Furthermore, a 11-item Research Preparation scale was developed to assess whether interns perceived they improved their understanding of the research process as a result of the program. A post-survey of mentors was similarly administered. Surveys were collected and managed using REDCap electronic data capture tools. REDCap (Research Electronic Data Capture) is a secure, web-based application designed to support data collection for research [23].

At the beginning of the vSREC, interns were invited to complete a pre-survey that collected demographic information, and a Research Preparation scale that evaluated the degree to which interns felt prepared and able to conduct research. Further, using free responses, interns were asked what they hoped to learn from the program and to discuss any concerns about the virtual research experience.

Interns completed a post-survey at the end of the program that included the same 11-item Research Preparation scale as well as scales evaluating the overall research and educational experience and evaluation of the research mentor. Free responses collected data about what interns learned from the research experience, future career goals, and their overall perception of the virtual research experience.

A Wilcoxon Signed Rank Test was used to compare interns’ responses on the Research Preparation scale, using SPSS v.27 (IBM Corp, Armonk NY). Significance was set at p ≤ 0.05. Responses to the post-survey were analyzed using descriptive statistics. Finally, free response data from pre- and post-surveys were coded and analyzed using thematic analysis.

## Results

### Pre-Program Concerns

All 22 interns in the vSREC completed the pre-survey. In a free-text response, six interns expressed concerns about having a different research experience due to the virtual format.

Specifically, they worried about not getting hands-on experience and having difficulty working with their mentors at a distance.

### Program Outcomes: Interns

A total of 18 interns completed the post-survey (12 in the SRP and six in the FSP, 82% response rate). All interns agreed or strongly agreed that their mentors were available to answer questions, provide advice, feedback, and provide resources to complete their research project (Figure 3A). Interns further described how mentors supported them in the summer research experience by making themselves available to answer questions, provide feedback, and offer mentorship beyond the program. Others described how their mentors were able to create a positive learning environment. One intern stated:

> *He created such a friendly and informative atmosphere. My mentor was very engaging and friendly during all our interactions, which definitely made me comfortable and content with my internship. In addition, he was able to explain very complex ideas in such a wonderful way! He started with the basics, then added fun anecdotes, until we could finally fully understand the more complex material. This helped to keep my interest level extremely high throughout all our interactions as well as during my independent study. His passion definitely rubbed off on me!*

**Fig. 3.**
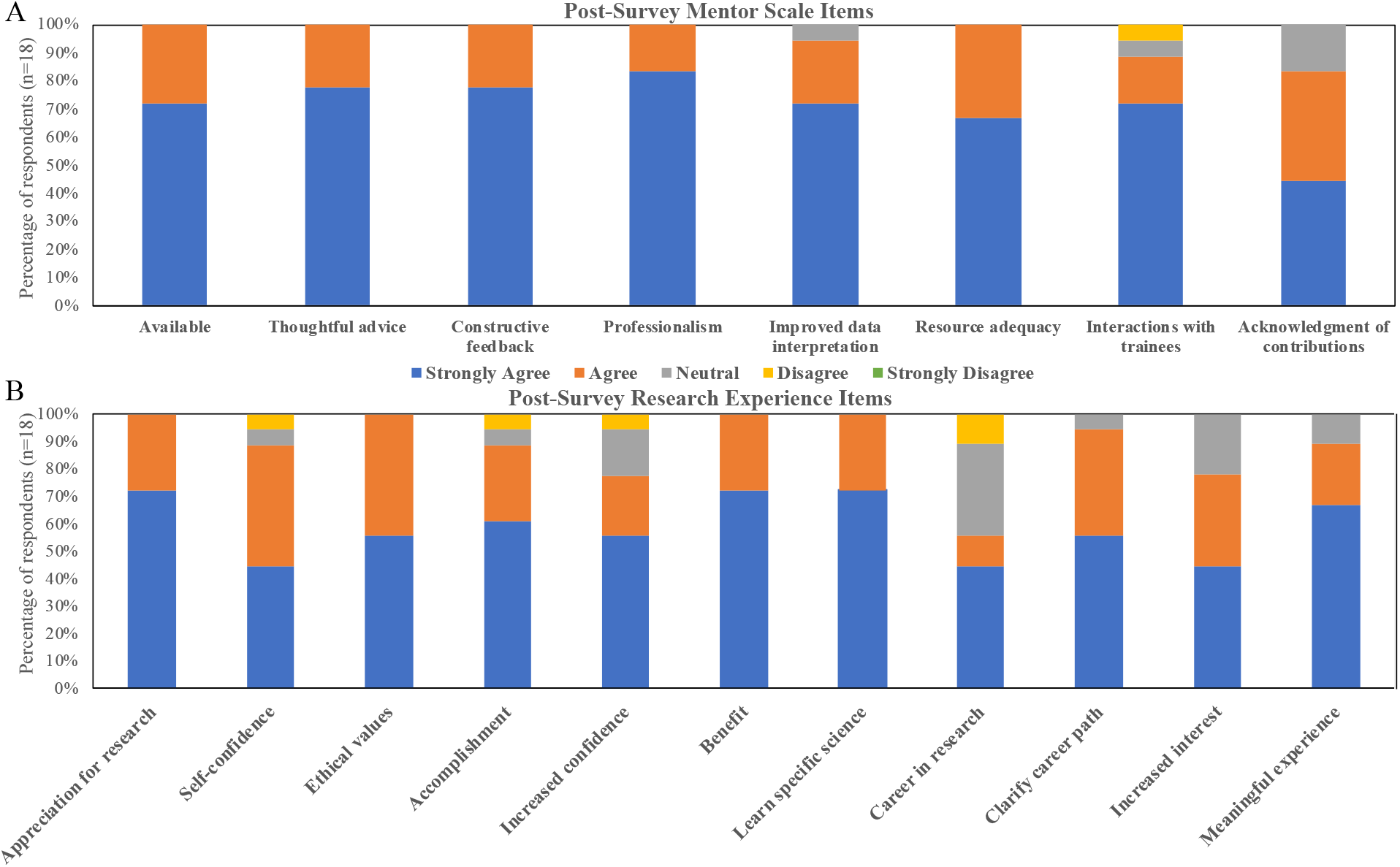
Evaluation of mentors and programs by vSREC interns. A) Evaluation of mentors by interns. B) The impact of vSREC on interns; results of post-program survey of n=18 interns.

Each intern rated the vSREC experience as good or excellent. All self-reported gaining a greater appreciation for research, learning ethical conduct of research, and studying a topic in depth. Fifty-five percent agreed or strongly agreed that they wanted to pursue a career in research (Figure 3B). Interns found expert speakers to be the most enjoyable aspects of the vSREC, followed by networking events. They appreciated the speakers sharing their research experiences and the pathways they took in their careers to reach their goals. One intern described:

> *The most enjoyable aspect of the virtual summer research experience was being able to hear all these different guest speakers who had such different paths that they followed to achieve their goal. It was overwhelming and very encouraging to continue chasing my dream after hearing all the bad experiences and setbacks that they experienced yet still managed to overcome.*

Interns also discussed the digital skills they learned during the virtual program and how they planned to use these in the future. Interns discussed how many of the skills in Zoom, Google Drive, and Canvas would assist them in college, particularly in online courses.

While the interns overwhelmingly enjoyed the virtual program, many discussed their challenges and recommendations for future virtual programs. Nearly half of respondents expressed a desire to have a hands-on research experience and felt it was difficult to sit in front of their computer for several hours a day. Interns recommended having more activities that were interactive to build rapport and engagement among interns and mentoring staff and to try to match interns in a lab with others at their level of education.

The Research Preparation Scale was included on both the pre- and post-surveys to evaluate interns’ perceptions of their ability to perform and conduct research after completing the virtual program. Interns reported a significantly greater understanding of the research process and in their preparation to conduct research. They also felt more able to analyze and interpret data and improved skills in scientific writing. Lastly, interns reported a greater understanding of how scientists conduct research, apply science to research, and apply the scientific method (Figure 4).

**Fig. 4.**
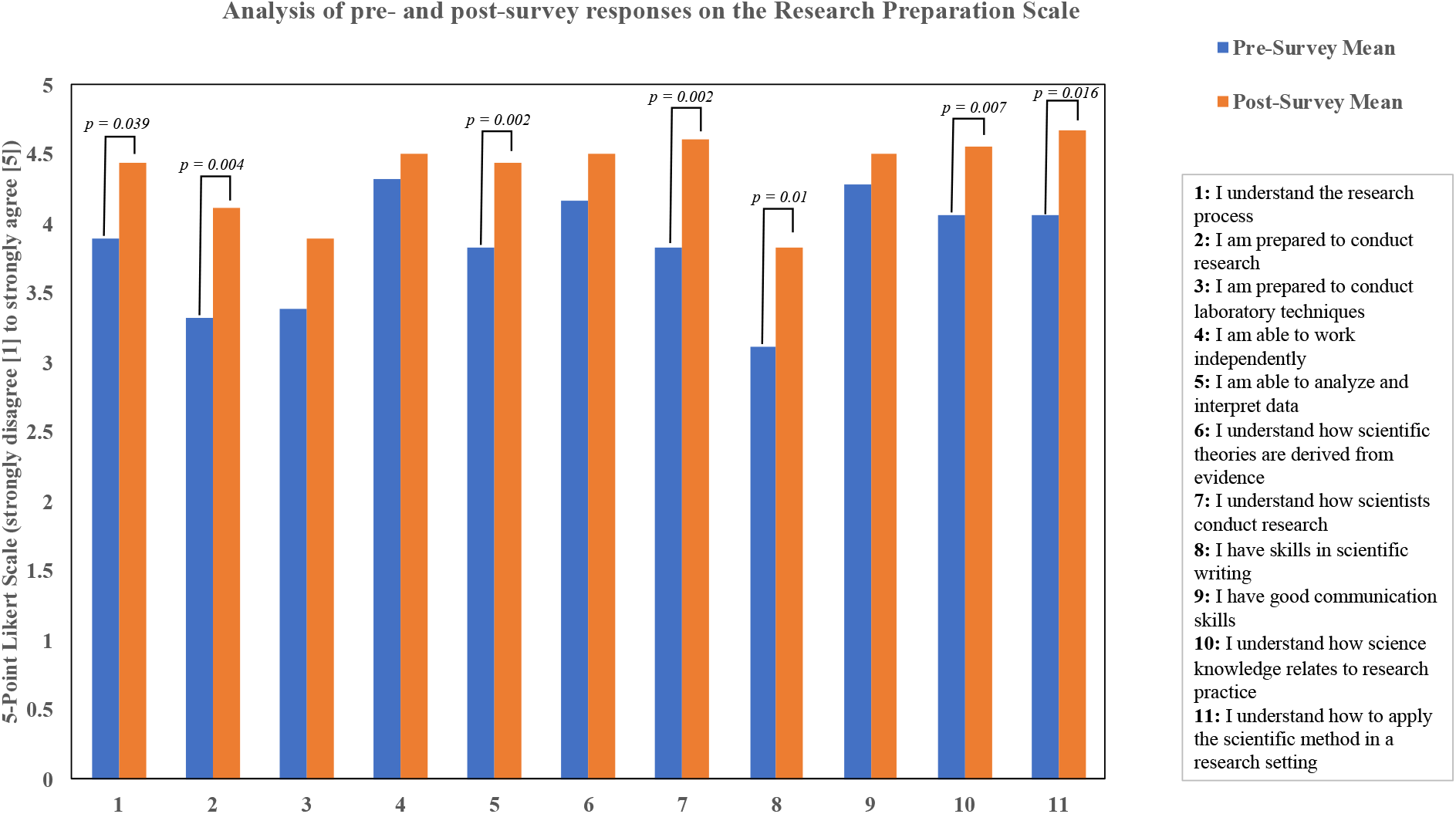
Comparative analysis of pre- and post-program survey results of interns’ pre-vSREC expectations and experience of vSREC. n=18 interns.

### Program Outcomes as Assessed by Mentors

At the end of the program, mentors were also sent a REDCap survey to assess their opinion of their interns’ progress. Of 10 respondents (out of 17 mentors), all agreed or strongly agreed (on a 5-point Likert scale) that “my intern gained understanding of how scientists work on real problems,” while 80% agreed or strongly agreed that “my intern learned digital research techniques” and “asked appropriate questions” (Figure 5). No mentors felt that interns spent too much time on other program activities, and 90% agreed/strongly agreed that program requirements enhanced the experience. However, 30% of mentors felt that interns were not on time or well prepared for Zoom meetings and commented anecdotally about varying levels of engagement and lack of clarity of expectations for both mentors and interns.

**Fig. 5.**
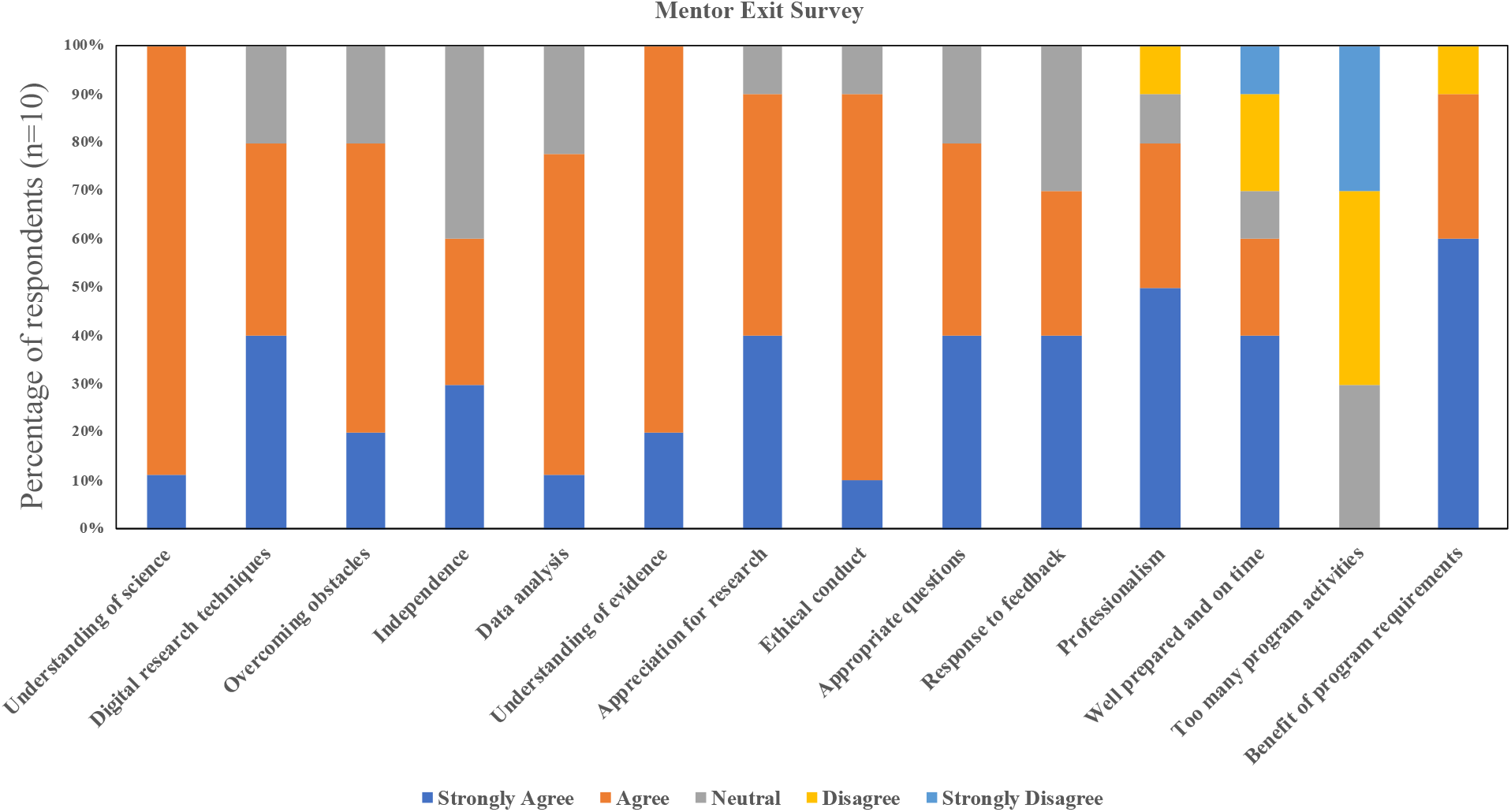
Evaluation of vSREC interns by mentors; results of post-program survey of n=10 mentors.

## Discussion

### Program Outcomes: Summary

As the COVID-19 pandemic unfolded in the USA, leading to lab hibernation in late March 2020, IUSCCC program leaders prioritized creation of these alternatives to hands-on research experiences. Due to stress associated with COVID-19, the abrupt closure of schools, and the resulting loss of social networks, the leadership of vSREC had tepid expectations for the program and expected student attrition. However, all but one student selected for the program based on an interview before hibernation readily accepted the offer to participate in vSREC. Moreover, each intern finished the program. The self-reported student and mentor outcomes strongly suggest a high degree of satisfaction with the program. However, there is an opportunity for further improvements, which are described below.

### Lessons Learned: Mentors and Interns

The research mentors provided a crucial link to research projects, models of cancer research career paths, and discipline-specific lecture topics, ensuring a cancer research focus was maintained. In the future, we envision additional innovative virtual projects in the areas of bioinformatics, image analysis, literature searches, and other *in silico* lab topics.

For interns, future plans might focus on professional communication. This training would include how to create a calendar-based schedule, how to schedule meetings on a mentor’s calendar, and professional etiquette for timeliness. Expectations related to intern-mentor interactions during the course of the program may further improve program experience, mentor satisfaction, and outcomes.

### Lessons Learned: Program Implementation

Active participation of high school teachers was key to the success of this program, as they applied their teaching and student-teacher interaction skills to keep interns engaged during the entire program. They also designed the curriculum shared by all interns, providing a common point of reference for all program participants. In the future, it will be valuable to draw on teacher expertise to design tests of student knowledge pre- and post-program, to ensure that self-reported learning achievements are supported by unbiased metrics.

Recruiting teachers for summer programs may pose a challenge in the future. Currently, teachers are seeing fewer opportunities for professional development within their schools because more time is being taken up to troubleshoot and prepare for the health and safety of the students within the virtual and in-person teaching platforms. This, in turn, creates fewer opportunities to enlist other strong teachers to help out with the summer program. A further challenge is that teachers are working longer hours to develop virtual and in-person lessons to accommodate the hybrid calendars created by most schools. This gives them less personal time to participate in professional development activities such as the Teacher Research Program. In the future, similar programs will need to consider innovative ways to recruit and retain strong teachers to help facilitate these high school pipeline programs.

The undergraduate students of the Summer Team were an invaluable part of the program, providing near-peer mentoring and technological support for online tools. The availability of computing devices and a good internet connection is a limiting factor for any virtual program. In an ideal program, tablets with cellular data connections would be made available to interns who need them. Although using multiple learning management system (LMS) platforms allowed for more comprehensive functionality than opting for a single standalone platform, this multi-platform use caused confusion for the interns, as they often struggled to remember the purpose of each platform. However, the benefits of this multi-platform method included access to the different native tools within each platform. No one platform provides all of the features needed to run a wholly virtual program. Still, training on integrating external platforms such as G Suite and Zoom into a central LMS system such as Canvas can help reduce some of the confusion that interns faced during the vSREC experience. Also, more comprehensive pre-program IT training for all interns by a member of the institution’s educational IT support team could help better prepare students for the upcoming program.

### Future Directions

A current problem in education is how to engage students with limited mobility (i.e., long-term wheel-chair bound or temporary injury limited mobility students) in the laboratory [24]. While the rehabilitation field has used adaptive sports as therapy [25], the adaptation of equipment and research facilities has been less swift. The opportunities for virtual research projects, such as those examples described here, offer a chance for meaningful engagement in cancer research to interns previously hindered by limited mobility. As work from home and telehealth becomes more accepted, we envision innovative opportunities to increase.

## Conclusions

The IUSCCC SRP program typically gets >200 applications for 15-17 slots. Thus, many students with interest in cancer research do not get the opportunity to participate. Further, IUSCCC summer programs do not provide a residential option, so many students from rural communities may be disadvantaged from participating. The virtual programs, however, offer the opportunity to engage students beyond geographic proximity to National Cancer Institute-designated cancer centers, particularly for those cancer centers that have entire states as their catchment area. Thus, a program such as this, developed in response to COVID-19, can potentially change the depth and breadth of cancer education. These impactful programs allow cancer centers to engage with communities. Although we hope that IUSCCC will be in a position to offer a hands-on laboratory experience in 2021, our virtual framework provides an appealing and effective alternative if needed.

## Supplementary Information

**Additional file 1.** NIH Definitions of students and underrepresented populations in science and from disadvantaged backgrounds. docx

**Additional file 2.** Detailed Program Design. docx

**Additional file 3.** Tables presenting vSREC final program, daily checkout survey, and Cancer 201 questions. xlsx

### Abbreviations

COVID-19: coronavirus disease 2019
FSP: Future Scientist Program
IUSCCC: Indiana University Simon Comprehensive Cancer Cancer
LMS: learning management system
NCI: National Cancer Institute
REDCap: Research Electronic Data Capture
SRP: Summer Research Program
TRP: Teacher Research Program
vSREC: Virtual Summer Research Experience in Cancer

## Declarations

### Ethics approval and consent to participate

This study received approval from the Institutional Review Board at Indiana University (IRB protocol #1110007280, approved 05/29/2020)

### Consent for publication

Not applicable.

### Availability of data and materials

The datasets used and/or analysed during the current study are available from the corresponding author on reasonable request.

### Authors’ contributions

TWC and SMH co-directed the program and contributed to the manuscript; ES managed the program and contributed to the manuscript; JB analyzed data and contributed to the manuscript; L-AC, JO, and ES were teachers in the Summer Team and contributed to the manuscript; RS, AB, OO, and JD were students in the Summer Team and contributed to the manuscript; AH and LC were teachers in the Summer Team; VB managed the program; HN oversaw the program and contributed to the manuscript. All authors read and approved the final manuscript.

## Acknowledgments

The authors wish to thank the leadership of IUSCCC and the Indiana CTSI for making financial and other resources available for effective implementation of the program. The authors also thank mentors and trainees and staff members of their labs for mentoring interns and faculty members of IUSM for didactic lectures. The authors thank Kaitlin Condron for the creation of display figures and IUSCCC for education program support.

