## Additional file 1 for "Building a virtual summer research experience in cancer for high school and early undergraduate students: lessons from the COVID-19 pandemic"

| **Additional file 1** NIH Definitions of students and underrepresented populations in science and from disadvantaged backgrounds. | | | |
| --- | --- | --- | --- |
| *To qualify for the program, a student must meet criteria of ONE of these categories* | | | |
| ***Categories*** | | **As defined by** | **Examples of groups** |
| **Racial and ethnic groups** |  | National Science Foundation | Black, African-American, Hispanic, Latinos, American Indian, Alaskan Native, Native Hawaiian, Other Pacific Islanders |
| **Individuals with disabilities** |  | Americans with Disabilities Act | Visual, hearing, walking, lifting, or cognitive disabilities |
| *Or, a student must meet criteria of TWO subcategories to qualify for the program.* | | | |
| **Disadvantaged backgrounds** | Homelessness | McKinney-Vento Homeless Assistance Act |  |
|  | Foster system | Administration for Children and Families |  |
|  | Eligible for free or reduced lunch | US Department of Agriculture |  |
|  | No parents with Bachelor’s degree | ED.gov |  |
|  | Eligible for Pell grants | ED.gov |  |
|  | Grew up in rural or low income area | Health Resources and Services Administration Rural Health Grants Eligibility Analyzer or Center for Medicare and Medicaid Services-designated low income and health professional shortage area |  |
