## Additional file 2 for "Building a virtual summer research experience in cancer for high school and early undergraduate students: lessons from the COVID-19 pandemic"

**Additional file 2: Detailed Program Design**

**A Summer Team to Support Students**

A “Summer Team” was created with the Program Directors, undergraduate students, and high school teachers from Indianapolis school districts to support interns in their research experience and to provide training on a core set of skills. Although there were overlapping areas of service, for simplicity, the undergraduate students helped with topics related to technology, and teachers supported topics related to the core curriculum.

**Harnessing Undergraduates’ Expertise and Learning Safe Use of Technology**

To effectively utilize numerous online tools and provide near-peer interactions, undergraduate students of the Summer Team provided training to vSREC interns on the Canvas Learning Management System (LMS), Google Drive, the full menu of options offered in Zoom, and the full functionalities of Exchange and Google email systems including rules, settings, calendar functions and proper email usage and etiquette. Because 9 of 22 (~41%) research interns were minors (age <18 years old), attention to online safety precautions was crucial. These precautions included instruction on Zoom etiquette, such as being present professionally on screen, selecting a private study area, use of headphones, and notification of family to avoid inadvertent background appearances. ‘Zoom-bombing’ was avoided by implementing waiting rooms, host controls, and registered participant lists for meetings. Total screen time was calculated and limited to five hours per day. Additionally, screen breaks were included into all didactics with an average of one 15-minute break for every two hours of screen time. Interns completed a daily checkout survey to summarize the events of their day, including breaks and stressors (see below). This alerted the Summer Team to any issues that interns were facing in order to individually address them during daily help sessions.

While vSREC interns predominantly used Zoom, Canvas, REDCap, and Google Drive during their summer experience, there are a plethora of alternatives to these systems. Future programs could use an alternative videoconferencing platform such as Microsoft Teams, Cisco Webex, or Google Hangouts based on availability of institutional licenses. Similarly, many LMS alternatives to Canvas exist as well, including but not limited to: Blackboard, Google Classroom, and Edmodo. Canvas was used for the vSREC experience due to Indiana University’s existing license and its seamless integration to other services such as Kaltura (video uploads), G Suite, and TopHat. Program coordinators should discuss options with IT specialists to identify existing platforms within the institution.

**Harnessing Talent from the Teacher Research Program and Developing a Core Curriculum**

To provide training on a common set of skills, high school teachers were selected from former Teacher Research Program (TRP) participants, and their pedagogical skills were harnessed to create a core curriculum of topics related to the research processes, scientific literacy, ethics, and grade-level resources for academic and career advancement. When possible, the Summer Team used existing training modules available through the Collaborative Institutional Training Initiative (CITI) to provide training on the Responsible Conduct of Research for Biomedical, Social and Behavioral Research, Human Subjects Research, Good Laboratory Practice, and NIH Recombinant DNA Guidelines. Proprietary modules available through Indiana University for developing skills in Excel, PowerPoint, Word, and other skills were also used. For specialized training, the Summer Team used expertise of campus experts. For instance, faculty from the Laboratory Animal Research Center (LARC) provided training on the guidelines governing the use of animals in research and faculty from the IUSM Ruth Lilly Medical Library taught a session on using full-text journal articles via PubMed and other search tools, and creating ORCIDs for each participant.

Where existing resources did not exist, the Summer Team created resources that could be used for future programs. These included materials on developing skills in navigating the Common App for colleges, resume creation, plus an extensive list of scholarships, and a listing of other research opportunities for interns to consider in future summers. These resources are available from the authors on request.

**Teamwork: Summer Team Supporting All Program Research Interns**

All program research interns were assigned a specific teacher to serve as their Student Coordinator. The role of each Student Coordinator was to facilitate the summer experience and provide assistance when needed via 1:1 and small group check-in meetings once per week. Each student had a personalized weekly schedule, based on their research mentor’s schedule (see below). The Summer Team was available daily for 1:1 “office hour”-type meetings to further explore and review topics, plus advise on personal and technological challenges.

**Meet-the-Expert Speaker Series**

To further augment the research intern experience, we sought to provide didactic sessions offering a diversity of topics reflecting the depth and breadth of expertise of the IUSCCC faculty, while ensuring that key scientific concepts were covered (see Additional file 3). The program began with a “Cancer 101” interactive Zoom lecture given by the program directors, introducing core concepts in cancer genetics and therapeutic modalities. The subsequent 16 interactive sessions engaged other experts and offered a vibrant and full perspective on both clinical and translational aspects of research in cancer. A minimum of two experts per week spoke on topics related to cancer, giving interns a full-spectrum orientation into the diverse career opportunities, research horizons, and clinical applications. Key cancer topics covered included: breast cancer research and treatment, lung cancer, drug discovery, immunobiology and immunotherapy, cancer prevention and control, and pediatric cancer. In addition, others spoke on their areas of interest, such as cancer virology, highlighting not just basic concepts but also current research. Faculty presenters modeled various career paths in cancer research and treatment, from PhD laboratory science faculty and population scientists, to nurses, to physician oncologists and surgeons. Finally, via Zoom, interns met a patient with ovarian cancer and her gynecologic oncologist. The patient was able to relate her own experiences from discovery to treatments and remission. Importantly, she emphasized the importance of knowledge, understanding, education, and community outreach.

We also deemed it important to cover topics that were timely and may have an impact on research interns outside cancer research. One week focused on COVID-19, with topics related to the SARS-CoV2 virus structure, innate vs adaptive immunity, testing windows and possible vaccines, which attracted many parents as audience members alongside the research interns. The vSREC program also engaged talented speakers on topics such as science communication, guidelines for use of animals in research, and career preparation from various local experts in these fields. Because the programs focused on underrepresented populations in biomedical sciences, we also aimed to provide hands on training for dealing with diversity and inclusion. Thus, the Indiana University School of Medicine Office of Faculty Affairs, Professional Development and Diversity gave an interactive, multi-media Zoom presentation on microaggressions.

Finally, interns engaged with Clinical Experience Day, a morning-long symposium that represented a spectrum of information on clinical trials. The day began with an interactive lecture on clinical trials, what they are, why we have them, and why they have regulations. Physician-scientists used this background information to describe the design, approval, and use of the human papillomavirus vaccine to prevent cervical and other cancers.

**Networking Events**

On Fridays, rather than having a didactic lecture, to finish up the week and create a well-rounded, holistic experience, interns had ‘Social Fridays,’ interactive networking events in which they got to learn important career tips and skills in a more relaxed, informal setting. This offered them an opportunity to learn about their peers’ research projects and to meet undergraduate students, graduate students, MD and MD-PhD students who each provided helpful advice on how to apply effectively for these programs and maximize each experience. For example, sessions were designed to provide tips for incoming college freshmen, others were for how to search and apply for scholarship opportunities, and others were to prepare the current undergraduates interested in medical school, graduate school or both. Eight such sessions provided interns a breadth of knowledge on various career paths, vertical mentoring and life of biomedical scientist (see Additional file 3).

**Mentor-Intern Integration Within the vSREC**

A consistent feature of our previous years’ programs was that the interns have always had access to a number of highly qualified IUSCCC mentors. Mentors are the principal investigator (PI) of a lab group and for vSREC were expected to meet with interns for at least two contacts per week. Contacts occurred in the form of 1:1 Zoom meetings, attendance at lab or group meetings, or meetings with multiple interns together. Additionally, each intern became a member of the lab group.

Interns completed several assignments from their lab groups, including interviewing a lab member, giving a journal club presentation, leading group discussion, and completing a virtual or *in silico* research project. The goal for interviewing a lab member was to learn about the educational background of the lab member that led to the current position and to learn a particular scientific technique in detail. In some lab groups, interns were paired and did journal club presentations on contemporary scientific literature. These assignments allowed vertical mentoring and built peer relationships. To increase the social and networking aspects of virtual integration, some lab groups had assignments of popular movies (i.e., *GATTACA*, *The Immortal Life of Henrietta Lacks*, *The Emperor of all Maladies*) followed by group discussion. These lab assignments were collected as part of the daily check in (see below).

The depth and breadth of research projects was vast. Some interns were given *in silico* data analysis projects, or projects that did not require generation of new data but required further analysis. Several interns used online databases to identify proteins with a motif found in the SARS-CoV2 novel coronavirus for potential therapy. Other interns were given electronic datasets for organization and analysis. Furthermore, several interns analyzed immunohistochemistry data using online quantification software. Some interns undertook literature reviews and discussion with PIs on biological and non-biological factors linked to cancer health disparity.

**Daily Check Out Evaluation**

Each day, interns completed a brief REDCap survey (see below) about the structure of their workday and what they learned from online modules, expert speakers, and mentor meetings. These daily checkouts served as a way for the Summer Team to ensure that interns maintained a healthy work/life balance and restricted program activities to “business hours”. Such a balance was essential to maintain in a virtual setting because, unlike in-person sessions, there are no concrete delineations of the start and end of the work day. An undergraduate student and a teacher from the Summer Team would both review the intern responses individually, then discuss which (if any) interns required follow up from a Summer Team member.

In the final week of the program, the program directors offered a “Cancer 201” question-and-answer session. The prompt for the questions was included in the Daily Evaluation the days prior. The questions asked by the interns in advance of this session highlight the high level of engagement, thought, and interest on diverse topics amongst the interns. Additional file 3 lists the types of questions put forward by interns and provides an example of how the program impacted scientific thinking in interns. Program directors categorized the questions, and used dice to randomly choose questions to answer during an interactive session.

**Institutional Commitment and Collaborations**

As noted, vSREC was made possible by the efforts of a diverse group of organizers. This extended to institutional support of the program. Our programs 2003 to 2019 were funded partially by a CURE supplement from the US National Cancer Institute, and through philanthropic support. Financial support for vSREC in 2020, which covered stipend and nominal financial lab support to mentors, came from philanthropic support to IUSCCC. To utilize existing resources on our campus, IUSCCC collaborated with the NIH Clinical and Translational Sciences Award-funded Indiana Clinical and Translational Sciences Institute (CTSI) to leverage its experience in education outreach, technology platforms, and relationships with the statewide network of all major academic and industry research centers. It was a mutually beneficial partnership as many of the resources created for the vSREC were also utilized by CTSI undergraduate and high school STEM outreach programs, and students in their programs were also invited to attend presentations hosted by IUSCCC faculty. Didactic lectures often had >100 attendees. CTSI infrastructure was also crucial for vSREC evaluation.
